# Conformational features and interaction mechanisms of VHH antibodies with β-hairpin-like CDR-H3: A case of Nb8-HigB2 interaction

**DOI:** 10.1101/2023.07.02.547379

**Authors:** Koichi Yamamoto, Satoru Nagatoishi, Makoto Nakakido, Daisuke Kuroda, Kouhei Tsumoto

**Author notes:** Correspondence and requests for materials should be addressed to S.N. and K.T.

## Abstract

β-hairpin conformation is regarded as an important basic motif to form and regulate protein-protein interactions. Single-domain V_H_H antibodies are potential therapeutic and diagnostic tools, and the third complementarity-determining regions of the heavy chains (CDR-H3s) of these antibodies are critical for antigen recognition. Although the sequences and conformations of the CDR-H3s are diverse, CDR-H3s sometimes adopt β-hairpin-like conformations. However, characteristic features and interaction mechanisms of β-hairpin-like CDR-H3s remain to be fully elucidated. In this study, we investigated the molecular recognition of the anti-HigB2 V_H_H antibody Nb8, which has a CDR-H3 that forms a β-hairpin-like conformation. The interaction was analyzed by evaluation of alanine-scanning mutants, molecular dynamics simulations, and hydrogen/deuterium exchange mass spectrometry. These experiments demonstrated that positions 93 and 94 (Chothia numbering) in framework region 3, which is right outside CDR-H3 by definition, play pivotal roles in maintaining structural stability and binding properties of Nb8. These findings will facilitate design and optimization of single-domain antibodies.

## Abbreviations

CDR, complementarity-determining region; MD, molecular dynamics; HDX-MS, hydrogen/deuterium exchange mass spectrometry; FR3, framework region 3; CD, circular dichroism; SPR, surface plasmon resonance; DSC, differential scanning calorimetry

## Introduction

The β-hairpin motif is a common protein secondary structure that often contributes to protein-protein interactions. In previous studies, cyclic peptides that adopt β-hairpin conformations have been shown to recognize multiple epitopes and regulate the activities of the proteins that they bind ^1–3^. Antibodies recognize various antigens with high affinity and selectivity and have been widely applied as diagnostic and therapeutic drugs ^4–6^. The complementarity-determining regions (CDRs) of antibodies play key roles in antigen recognition and often form β-hairpin-like structures ^7, 8^. Therefore, β-hairpin would be one of the significant basic motifs to adjust the binding and activity regulation for proteins.

Recently, single-domain V_H_H antibodies derived from the N-terminal domain of camelid heavy-chain antibodies have received considerable attention ^9–11^. V_H_H antibodies have multiple advantages compared to conventional IgG or Fab antibodies including lower molecular weight (∼15 kDa), better cell permeability ^12, 13^, and lower production costs ^9–11^. Moreover, V_H_H antibodies are better able to recognize clefts or concave pockets of antigens than are conventional IgG or Fab antibodies ^14–19^. In 2018, caplacizumab became the first clinically approved V_H_H antibody ^20^. Applications of V_H_H antibodies would be expected to be developed.

The third CDR (CDR-H3) of V_H_H antibodies is especially critical for antigen recognition ^17^. When the length of CDR-H3 is short, it often adopts a extended β-hairpin-like conformation, similar to that of conventional antibodies ^21–23^, but this conformation is less common than a helical-bending conformation ^22, 23^. Due to the rarity, few researchers analyzed the structural characteristics of β-hairpin-like CDR-H3 in detail. However, short CDR-H3 would be suitable to obtain high affinities against glutathione S-transferase (GST), which is widely utilized as a tag sequence for recombinant protein expressions ^24^.

Thus, there would be some possibilities for β-hairpin CDR-H3 to prefer to recognize unique epitopes, different from helical-bending CDR-H3s. We reasoned that if structural features of a β-hairpin CDR-H3 were elucidated, it would lead to the improvement of artificial V_H_H library designs and more accurate structural prediction. Therefore, we investigated the molecular mechanisms of recognition of the llama V_H_H antibody Nb8, which binds specifically to HigB2 and that harbors a β-hairpin-like CDR-H3 (PDB ID: 5mje ^25^). The molecular mechanisms were analyzed by alanine-mutation experiments, molecular dynamics (MD) simulations, and hydrogen/deuterium exchange mass spectrometry (HDX-MS). We identified structurally important residues in the framework region 3 (FR3). Our findings will be useful for molecular design of β-hairpin CDR-H3s.

## Results

### Investigating significant residues upon the antigen recognition among CDR-H3 and its surrounding amino acids using SPR (CDR-H3, FR3, FR2)

As a model V_H_H with an CDR-H3 that adopts a β-hairpin-like structure, we selected Nb8. A crystal structure of the complex of Nb8 with its antigen HigB2 has been solved (Figure 1, PDB ID: 5mje) ^25^. We identified the CDR-H3 region using SabPred-ANARCI ^26^, based on the Chothia numbering scheme ^27^ (Table 1). To determine whether the residue features were specific to V_H_H antibodies or not, the frequencies of each residue in llama V_H_H antibodies and human antibodies were analyzed using the AbYsis database ^28^. Buried surface area (BSA), accessible surface area (ASA), and polar interaction of each residue for the antigen recognition were examined using the PDBePISA server ^29^ (Table S1).

**Figure 1.**
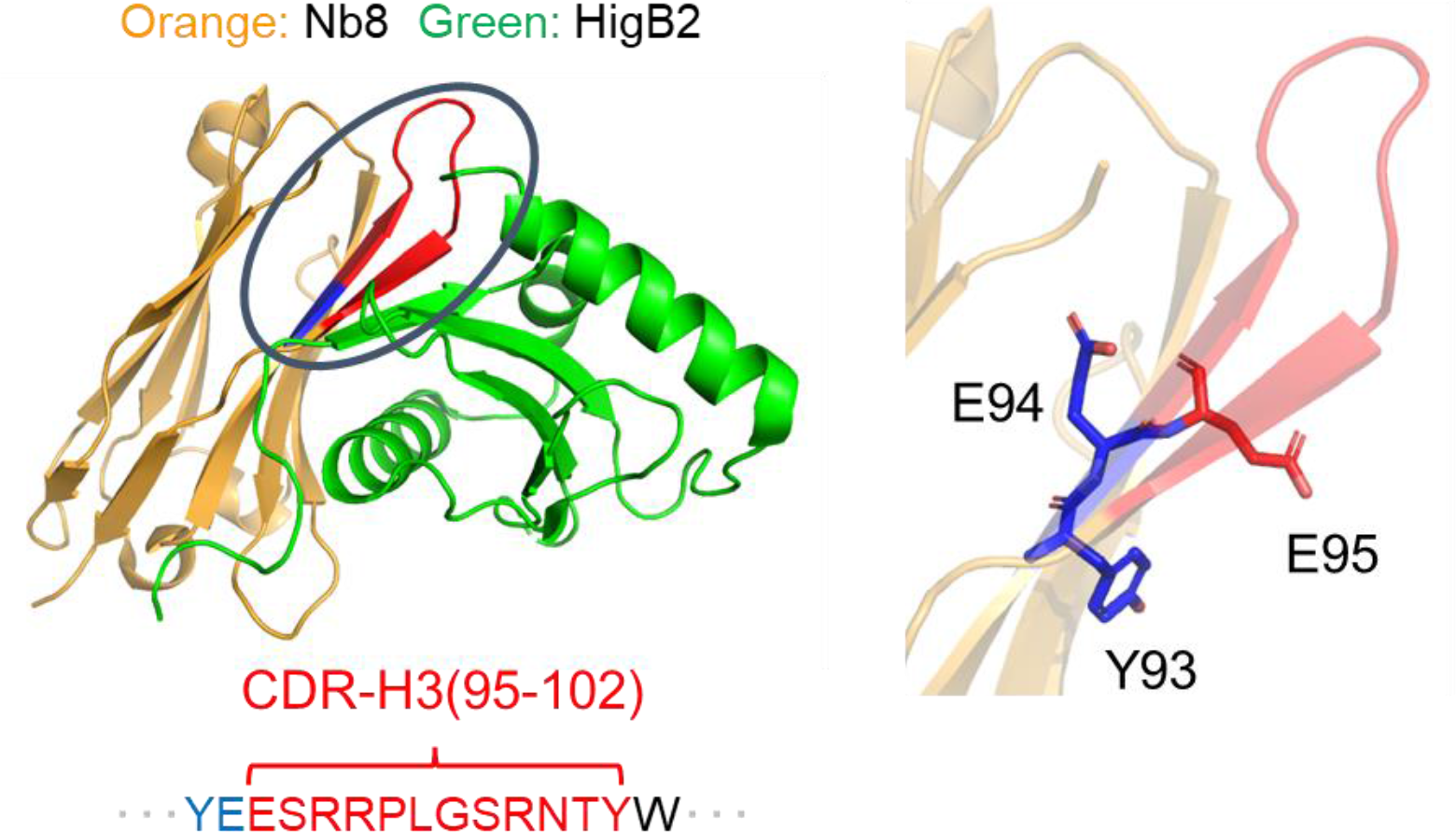
Crystal structure of model antibody-antigen complex of Nb8 with HigB2 (PDB ID: 5mje). Left: Ribbon diagram of the complex. With the exceptions of CDR-H3 (amino acids 95-102) and surrounding residues (amino acids 93, 94), which are indicated in red and blue, respectively, the antibody is depicted in orange. The β-hairpin-like region is circled. Right: Y93, E94, and E95 of Nb8 were depicted by sticks.

**Table 1.** CDR and FR regions of Nb8 antibody (Chothia numbering). Blue: residue 93, 94.

|  | **FR-H1**<br>**(1-25)** | **CDR-H1**<br>**(26-32)** | **FR-H2**<br>**(33-51)** | **CDR-H2**<br>**(52-56)** |
| --- | --- | --- | --- | --- |
| **Nb8** | QVQLQESGGGLVQP<br>GGSLRLSCVVS | GDRRTIY | TMGWFRQAPGNQGELV<br>ATM | TSSGV |

|  | **FR-H3**<br>**(57-94)** | **CDR-H3**<br>**(95-102)** | **FR-H4**<br>**(103-113)** |
| --- | --- | --- | --- |
| **Nb8** | TTYVDSVKGRFSISRDSAEDSAK<br>NTVSLQMNSLKPEDTAFYTCYE | ESRRPLGSRNTY | WGQGTQVTVSS |

To investigate the function of β-hairpin-like CDR-H3 in detail, alanine-scanning mutations were made in Nb8, and the affinities of mutants for HigB2 were determined. The target amino acids were mainly located on CDR3 and the surrounding regions, based on the antibody sequence and interaction information as shown in Table 1. Surface plasmon resonance (SPR) was used to determine dissociation constants (*K*_D_) and kinetic parameters (*k*_on_, *k*_off_) (Figure 2, Figure S1, Table S2). When mutations were made in the CDR-H3 region, drastic declines in binding affinities were observed. Mutation of either R97 or R100c of Nb8 considerably reduced its affinity for HigB2 (WT: 17.0 ± 0.04 nM, R97A: 3.02 ± 0.09 µM, R100cA: 5.50 ± 0.04 µM). In FR3, residues Y93 and E94 were critical (Y93A: >16 µM, E94A: 5.36 ± 0.12 µM). In FR4, mutation of W103 considerably reduced the affinity for HigB2 (W103A: 1.27 ± 0.02 µM). All of these changes resulted from large increases in *k*_off_ parameters. The dissociation rates were too high to calculate kinetic parameters for the Y93A mutant. Other kinetic features in CDR-H3, even though no apparent shift in binding affinity was observed upon mutation of E95, a gain in *k*_on_ value was detected.

**Figure 2.**
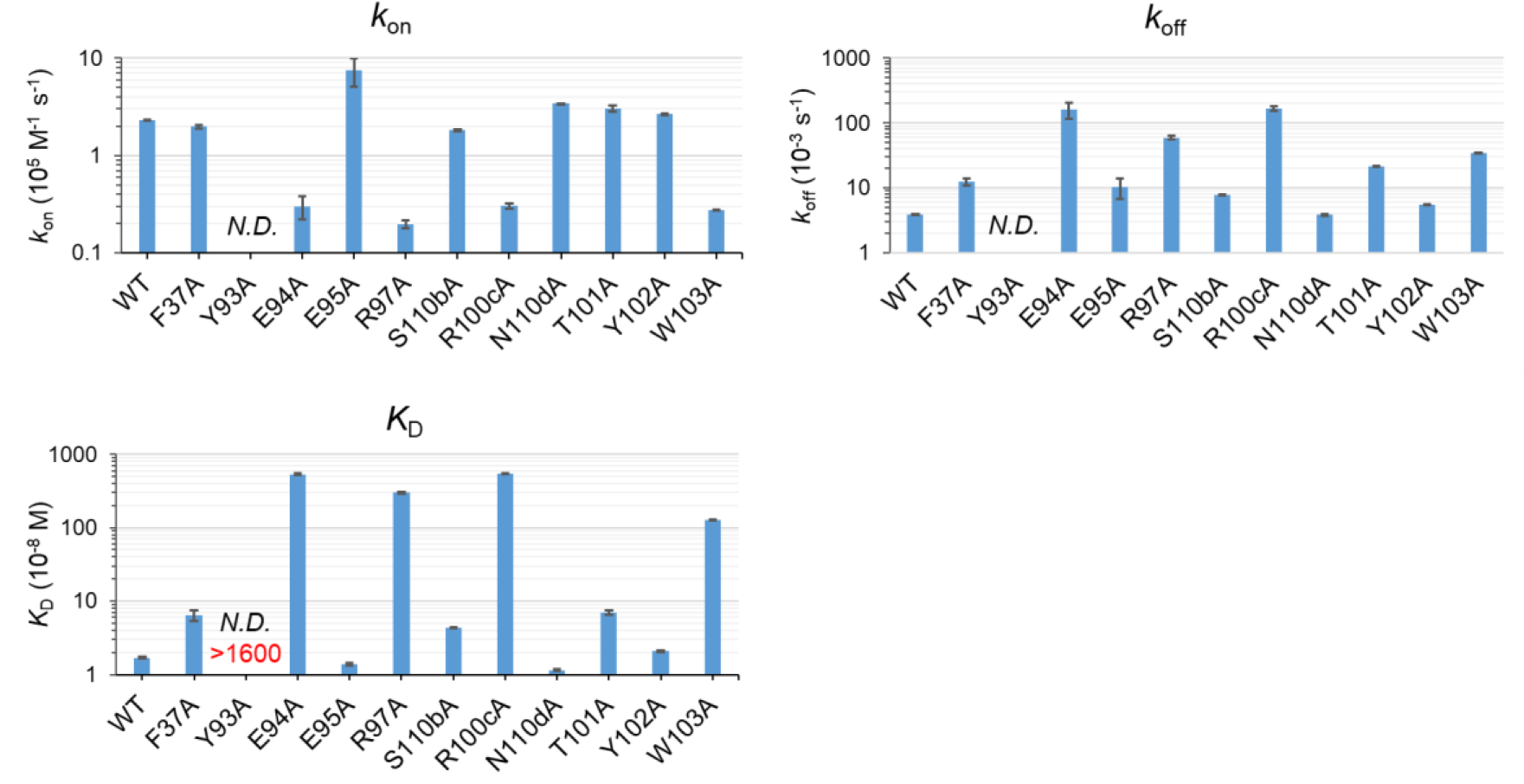
Kinetic parameters of wild-type Nb8 and mutants determined using SPR. Error bars correspond to the standard deviation. Values were determined at 25 °C. *N.D.*: not determined.

### Interaction analysis of hot spot residues in CDR-H3 upon the antigen (R97, R100c)

In the co-crystal structure, R97 of Nb8 forms hydrogen bonds and salt bridges with E12 of HigB2. Neither the thermostability (Table 2, Figure S2) nor the secondary structure (Figure S3) of the R97A mutant differed from those of the wild-type Nb8, suggesting that R97 would be the hotspot residue for antigen recognition. The side chain of R100c forms hydrogen bonds and salt bridges with E7 of HigB2, and the main chain of R100c forms a hydrogen bond with HigB2 E12. Even though the thermostability of R100cA was decreased relative to the wild-type Nb8 (Table 2, Figure S2), the secondary structure was maintained (Figure S3). Therefore, R100c is also likely critical for antigen recognition.

**Table 2.**
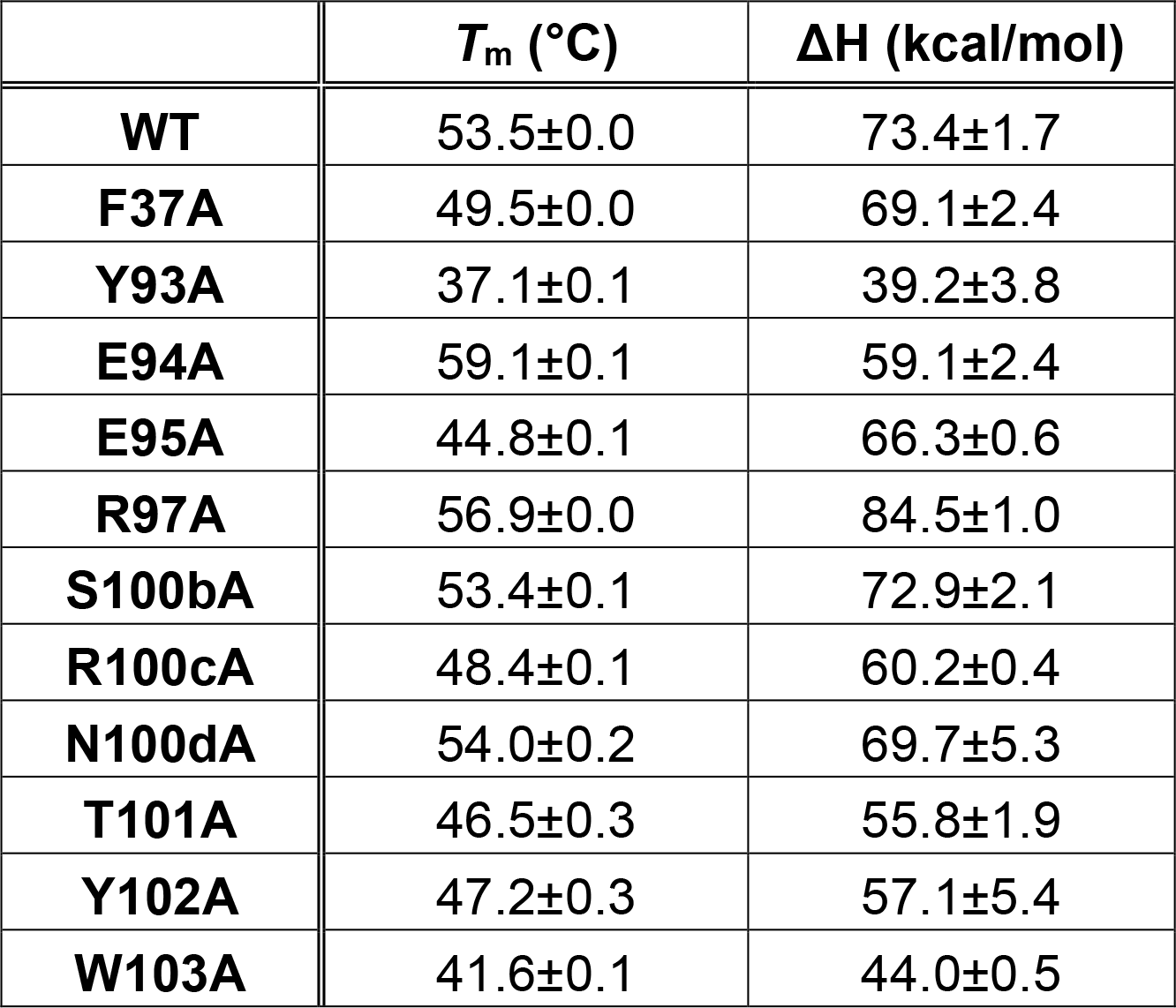
Thermostability parameters of wild-type Nb8 and mutants determined using DSC. Means and standard deviations are given.

To determine whether there are intramolecular interactions between R97 and R100c, we conducted MD simulations on the apo-form of Nb8. In this simulation, R97 and R100c were in close proximity (Figure S4), and the distances between these two positive charges were less than or equal to 10.0 Å in 51.1% of the simulations. Charge-charge interactions can be generated even when distances are between 6-10 Å ^30^. Thus, these data suggest that there is electrostatic repulsion between R97 and R100c. Differential scanning calorimetry (DSC) measurements indicated that the R97A mutant was more stable than the wild-type Nb8 (Figure S2). This result confirms that there is electrostatic repulsion between R97 and R100c.

### Structurally important roles of W103 in FR4

W103 of FR4 is highly conserved in V_H_H antibodies and conventional antibodies (92.9% identity in llama V_H_Hs, 99.2% in human antibodies based on the abYsis database). In conventional antibodies, W103 is located at the VL-VH interface, and it does not directly interact with antigen. Previous reports indicate that W103 is important for the structural stability of V_H_H antibodies ^31–33^. In Nb8, W103 main chain atoms form two hydrogen bonds with S3 of the antigen. More strikingly, we observed an apparent loss of thermostability when W103 was mutated to Ala: The melting temperature (*T*_m_) of the mutant was 11.9 °C lower than that of the wild-type protein (Table 2, Figure S2). Significant changes in secondary structures were also detected by CD analysis (Figure S3).

### Interaction analysis of Y93 in FR3

The buried surface area (BSA) of Y93 was not as large (22.4 Å^2^) as those of other residues of Nb8 in the complex with antigen (Table S1). However, Y93 forms a cation-π interaction with the side chain of R25 in HigB2, and the ratio of BSA from accessible surface area (ASA) was relatively large (90.0 %). The thermostability of the Y93A mutant was very low (*T*_m_ 37.1 °C, Table 2, Figure S2), and the CD spectrum was distinct from that of the wild-type protein (Figure S3). These phenomena showed the significance of Y93 on the folding of the V_H_H antibody. To comprehend the function of Y93, we examined intramolecular interactions of Y93 using MD simulations. In 80.3% of simulations, Y93 was close to W103 with centroid-centroid distances less than or equal to 6.50 Å (Figure 3A, Figure S5). This close contact was observed the crystal structure of the Nb8 complex with HigB2 (centroid-centroid distance: 5.1 Å). In V_H_H antibodies, V37F/Y mutation from human antibodies is widely known ^19, 23, 31^, and the closely situated position of F/Y37 and W103 is observed ^33–36^. Also, when V37F substitution was applied to VH antibodies, side chain orientation of W103 was arranged to be able to create T-shaped π-π stacking with F37 ^34, 37^. Thus, this aromatic interaction likely contributes to the stability of V_H_H antibodies. In the co-crystal structure of Nb8, F37 and W103 are located at a centroid-centroid distance of 6.4 Å; however, in MD simulations, side chains of F37 and W103 were located at T-shaped geometry and the centroid-centroid distance was less than 6.50 Å in 85.9% of simulations, suggesting the presence of π-π stacking. Therefore, Y93 would interpose the F37-W103 interaction and highly contribute to conformational stabilization.

**Figure 3.**
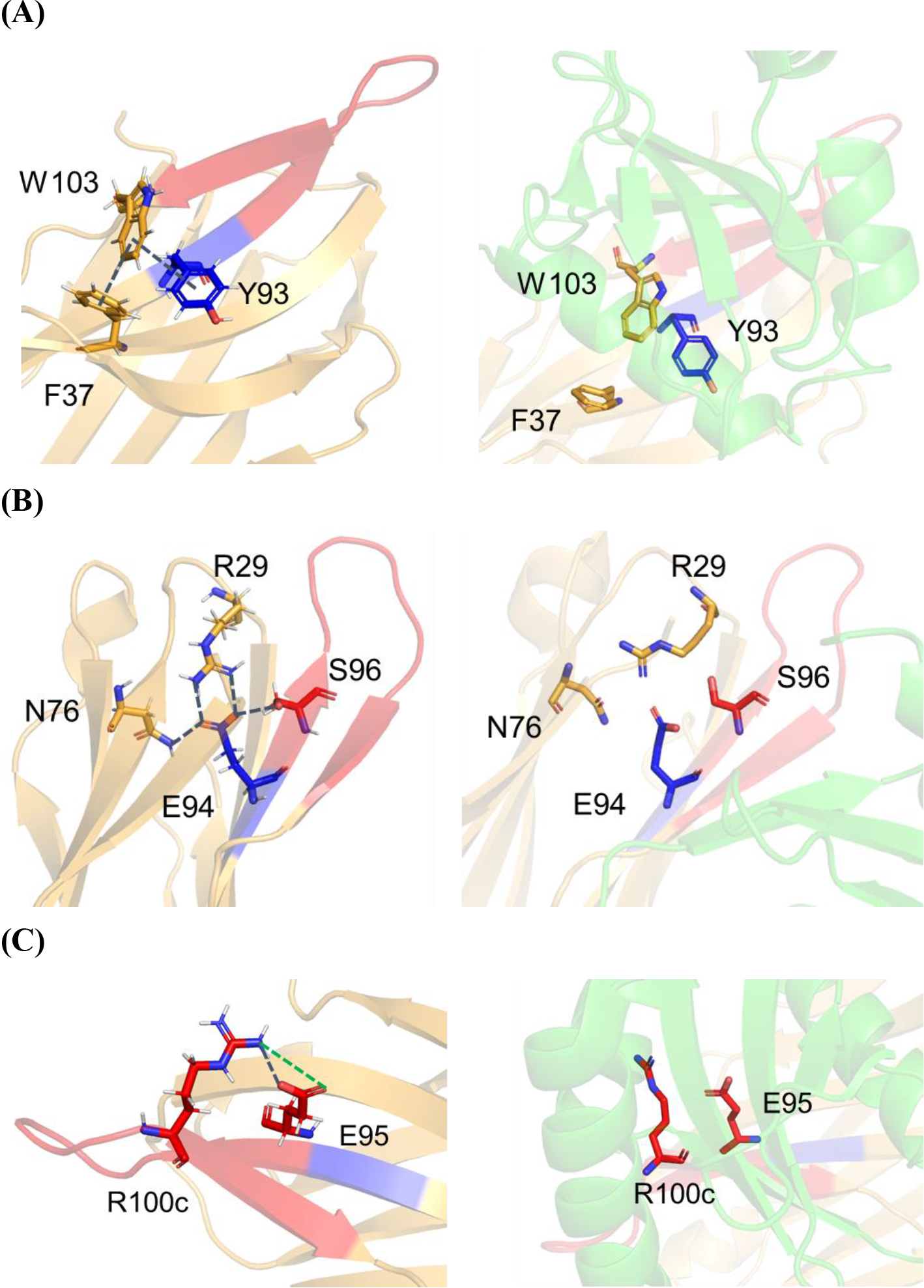
Positions of residues involved in intramolecular interaction networks in Nb8. In each panel, the structure from run 1 of the MD simulation (200.00 ns) is shown on the left and the co-crystal structure is shown on the right. (A) Y93-associated aromatic interactions. (B) The buried hydrogen bond network of E94. (C) Hydrogen bonds and salt bridges between E95 and R100c. Each interaction in the MD simulation is depicted by a dashed lines, and stability of each distance is shown in supplemental information.

### Interaction analysis of E94 in FR3

E94 in FR3 is almost totally buried inside Nb8, and no direct interactions are observed with the antigen. Interestingly, the *T*_m_ of the E94A mutant was 5.6 °C higher than that of the wild-type protein (Table 2, Figure S2). We also observed a drastic difference in the CD spectrum of E94A compared to that of the wild-type Nb8 (Figure S3). This indicated that E94 influences the folding of Nb8. In MD simulation, stable hydrogen bonds were formed between E94 and R29 in CDR-H1 (Figure 3B, Figure S6). Two hydrogen bonds between R29 NH2-E94 OE1 and between R29 NH1-E94 OE2 at distances less than 3.50 Å were detected in 69.2% and 79.5% of simulations, respectively. Simulations also indicated that Asn76 in FR3 and S96 in CDR-H3 interacted with E94 (Figure 3B, Figure S6). In co-crystal structure, only the interaction between E94 and R29 was detected.

### Interaction analysis of E95 in CDR-H3

Mutation of E95 in CDR-H3 had little influence on the binding affinity, even though it has several direct interactions against the antigen. However, the kinetic parameters of E95A differ from those of the wild-type Nb8: E95A had a higher *k*_on_ value (Figure 2). In the co-crystal structure, E95 forms hydrogen bonds and salt bridges with R15 and R25 of HigB2. In MD simulations, E95 OE1 and R100c NH2 were within 4.00 Å or 3.50 Å in 59.5% or 48.0% of simulations. Also, E95 OE2 and R100c NH2 were within 4.00 Å or 3.50 Å in 59.6% or 48.5% of simulations (Figure 3C, Figure S7), indicating that salt bridges and hydrogen bonds were often formed between these residues. However, these intramolecular interactions were disrupted when Nb8 associated with HigB2. Similar to R100cA, the E95A mutation resulted in a loss of thermostability (Table 2, Figure S2), hence E95-R100c interaction would play a key role on retaining the stability of V_H_H antibody. On the other hand, this interaction should be disrupted upon antigen binding, thus E95 has a negative effect on *k*_on_ parameter.

### Folding dynamics for alanine mutants of Position 93, 94, and 95 in Nb8

We next performed hydrogen/deuterium exchange mass spectrometry (HDX-MS) analysis to evaluate the folding or conformational dynamics of Y93A, E94A, and E95A mutants and the wild-type Nb8 (Table S3). We found that the Y93A mutant was more likely to undergo hydrogen/deuterium exchange throughout its length than was the wild-type protein (Figure S8A). This result indicates that Y93A destabilizes the entire V_H_H structure. The E94A mutant had significantly increased exchange rate in the CDR-H1 region compared to the wild-type protein (Figure S8B, C). In the E95A mutant, the hydrogen/deuterium exchange rate was slightly increased compared to the wild-type protein in the CDR-H1 region as well as in the CDR-H3 and FR4 regions (Figure S8D, E).

## Discussion

We analyzed binding affinity to the antigen, melting temperature, secondary structure, and the folding dynamics of the wild-type and mutant antibodies. We identified several important amino acids in CDR-H3, FR3, and FR4. CDR-H3 is the CDR region primarily responsible for antigen binding, and the R97 and R100c in this region directly interact with the antigen HigB2. W103 of FR4 is a highly conserved amino acid in V_H_H antibodies that has been shown to be important for antibody stability and binding activity in other studies ^31–33^, and our study showed a similar function. Mutation of Y93 and E94, located in FR3, and E95 in H3 also caused significant changes in kinetic and thermal stability compared to the wild-type protein. These FR3 residues have not been discussed well in previous studies, and their importance may be characteristic of Nb8 antibodies. These three amino acids are discussed in detail below.

### The Role of Position 93 in Nb8: pivotal contribution for stabilizing antibody structure via aromatic interactions

In Y93A mutant, obvious damages on the structure of Nb8 were shown in CD and DSC analyses. During the preparation of this antibody, yields were low, and the mutant eluted during size exclusion chromatography at a different retention time than the wild-type protein (Figure S9). In HDX-MS analysis, Y93A was also observed to have a distorted overall V_H_H structure. Based on these data, we concluded that Y93 is critical for proper folding of Nb8. We checked the frequencies of amino acids in heavy chain position 93 using the abYsis software. Position 93 is the most often Ala prominent residue in V_H_H antibodies, and Asn is next most common (Figure 4A). In human antibodies, Ala is also the most common residue (Figure 4A). The portion of Y93 is not very high even among V_H_H antibodies, it has the fourth highest frequency (2.9 %) in llama V_H_H. Conversely, Y93 is very rare (<0.1 %) in human antibodies. Highly conserved residues often play significant roles in the structural stability and physical properties of antibodies ^7, 31, 33^. The importance of Y93 appears to be a unique feature of V_H_H antibodies.

**Figure 4.**
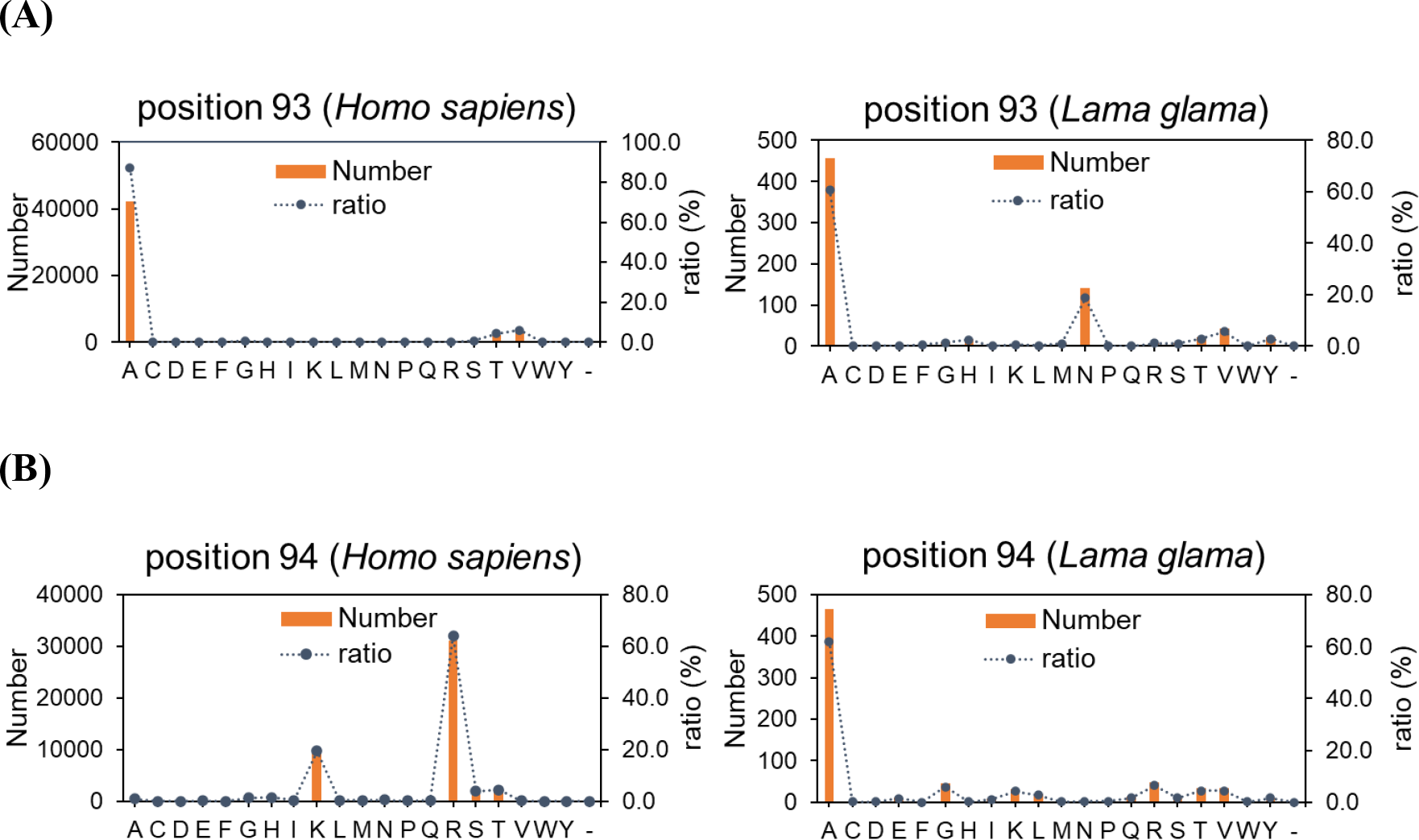
Amino acid frequencies in the heavy chain at (A) position 93 and (B) position 94 of in *Homo sapiens* and *Lama glama* antibodies. Data from abYsis database, accessed December 12, 2022.

In the MD simulation, Y93 and W103 were located in close proximity with centroid-centroid distances of less than or equal to 6.50 Å in 80.3% of simulations. In most V_H_H antibodies, W103 forms aromatic interactions with the residue at position 37, which is usually Phe or Tyr, that contribute to the conformational stability ^31, 33, 36^. As expected, the W103A mutant had a much lower *T*_m_ than the wild-type protein; however, the F37A mutant was only slightly less stable than the wild-type Nb8. Therefore, in our model system, the F37-W103 interaction was not critical. In contrast, disruption of Y93-W103 interaction disarranged not only the β-hairpin conformation of CDR-H3 but the overall fold of Nb8. Therefore, in Nb8 case, Y93 was purposely selected in order to make additional Y93-W103 interactions and maintain stable whole antibody structure.

### The Role of Position 94 in Nb8: conformational rearrangement via buried hydrogen bond network

Position 94 position is Ala in 62.1% of V_H_H antibodies. In human antibodies, this position is usually Lys or Arg residues (19.5% and 64.1%, respectively). Thus, the E at position 94 is a specific characteristic of Nb8 (Figure 4B). In Nb8, E94 forms intramolecular interactions with R29, regardless of the presence of the antigen. Considering the increase of thermostability on E94A mutant, E94 would improve antigen recognition in expense for stability loss. When R29, Asn76, or S96, which formed intramolecular interactions with E94 in MD simulations, were mutated, thermostability decreased (Δ*T*_m_ = −10.1 °C for R29A, −8.1 °C for S96A, Figure S10). HDX-MS analysis confirmed that the E94A mutant was more flexible than the wild-type protein. Therefore, the presence of E94 would be unfavorable for thermostability, and the stability loss was reduced by the surrounding residues. It is suggested that E94 creates a moderate flexibility to produce the antigen-binding activity. Δ*H* of E94A became small in DSC analysis, and E94 would adjust structural entropy. Therefore, E94, located in front of CDR-H3, is not involved in direct binding to the antigen, and would have the main role in maintaining the structure of CDR-H3 and the whole conformation of Nb8 to be appropriate for antigen recognition.

### The Role of Position 95 in Nb8: Supporting a hotspot residue on the basis of thermostability

Mutation of E95 in the CDR-H3 to Ala had almost no effect on secondary structure, but it did decrease thermostability. The binding affinity for antigen was similar to that of the wild-type Nb8 with a slight increase in *k*_on_ value. In MD simulations, E95 often formed hydrogen bonds with R100c, another critical residue in CDR-H3. HDX-MS analysis showed that the E95A mutant had increased the flexibility in CDR-H3 and FR4 regions compared to the wild-type protein. R100c is necessary for selective binding to HigB2, and E95 would adjust CDR-H3 conformation preferring for better thermostability.

### Role of FR3 residues (93, 94) in β-hairpin CDR-H3 of V_H_H

From searching for additional V_H_H antibody models harboring Y93 by SabDab-nano ^38^, we picked and examined co-crystal structures whose main chains of CDR-H3 were identified. Of the 10 structures identified, nine had CDR-H3s with extended β-hairpin-like conformations (Figure S11). When polypeptide chains form β-hairpin structures, paired cross-strand residues on opposite β-strands are either hydrogen-bonded or non-hydrogen-bonded positions ^1, 39^. If two aromatic residues are present at non-hydrogen-bonded positions, π-π stacking between the two side chains strongly stabilizes the β-hairpin conformation ^40–42^. In Nb8, Y93 and W103 are located at non-hydrogen-bonded positions. Tyr at position 93 is not common in V_H_H antibodies. A reference study for designing extended-type CDR-Hs3 diversified amino acids on position 93, and Asn was most frequently selected; however, Tyr was not included ^21^. In addition, we confirmed the BSA/ASA ratio of Y93 for the antigen binding, and high coverage (average: 76.4 %) was observed. In some V_H_Hs with Tyr at position 93 (e.g., PDB ID: 6itc_V, 7m1h_E), the side chain of Tyr forms hydrogen bonds with antigen. Hence Y93 would have additional roles contributing to direct interactions with the antigens. Thus, position 93 would be one of the significant residues on extended β-hairpin-like CDR-H3. For future research, position 93 would have high significance for designing V_H_H antibodies harboring extend-type CDR-H3.

In conventional human antibodies, heavy chain position 94 is usually either Lys or Arg (19.5% and 64.1%, respectively). Interactions such as intramolecular hydrogen bonds or a salt bridge are observed between position 94 and D101, and the residue types at these locations have been utilized to predict CDR-H3 conformations ^7^. This position influences affinity as shown by substitution from S94 to R94 ^43^. Thus, this position is important in conventional antibodies. In llama V_H_Hs, Ala is the most frequently observed amino acid at position 94 (62.1%), and conventional structural prediction criteria used for CDR-H3 regions of conventional antibodies are not as accurate for llama V_H_Hs ^23^. In Nb8, E94 is almost fully buried into the antibody (ASA = 1.34 Å^2^), and it forms internal hydrogen bond networks. Even though E94 was a specific feature of Nb8, the surrounding environment and structural contribution of position 94 would be different from conventional antibodies, and the high buried degree may also be one characteristic of β-hairpin-like CDR-H3 in VHH antibodies. Therefore, further research would be needed to check the validity and expansivity of the supposition.

A stable CDR-H3 conformation is likely important for both the thermal stability of and for antigen recognition by V_H_H antibodies ^44, 45^. The helical-bending conformation is the most common conformation of CDR-H3 regions of V_H_Hs, and the influences of mutations in FR-H2 on this conformation have been explored ^23, 31, 46^. For instance, V37F/Y, G44E, L45R, and W47F/G substitutions were the notable differences between V_H_H antibodies and conventional antibodies ^23, 31, 46, 47^. Mutation to human sequence decreases thermostabilities and affinities of V_H_H antibodies ^31, 36^. Interestingly, the FR residues we identified in Nb8 as important for antigen recognition were not at these positions. Our identification of FR3 amino acids that are critical to a V_H_H with a β-hairpin CDR-H3 will provide valuable insights for the future design of artificial V_H_H libraries and computational evaluation and engineering of V_H_Hs.

### Conclusion

The CDR-H3 regions of single-domain V_H_H antibodies sometimes form β-hairpin-like conformations that are critical for antigen recognition; however, specific features and interaction mechanisms of such β-hairpin-like CDR-H3s are not well understood. Using a model V_H_H antibody-antigen we identified positions important for thermostability of the antibody and for high-affinity antigen binding. Our study also revealed novel mechanisms of FR3 residues, which impact properties of the antibody. These residue features would be unique to β-hairpin-like CDR-H3s of V_H_H antibodies, and our findings will facilitate engineering of V_H_H antibodies.

## Materials and methods

### Cloning, expression, and purification of V_H_H antibodies

A synthetic gene encoding Nb8 was optimized for expression in *Escherichia coli* and cloned into the pRA2 vector ^48^. The construct contained a pelB signal peptide at the N-terminus and a His_6_-tag at the C-terminus. Mutants were generated by site-directed mutagenesis using a KOD-Plus Mutagenesis Kit (Toyobo, Japan) or PrimeSTAR® Mutagenesis Basal Kit (Takara Bio, Japan). For the expression, *E.coli* BL21(DE3) cells carrying the expression vector were grown in 1 L of LB medium containing 100 µg/mL ampicillin at 37 °C and 120 rpm. When the optical density at 600 nm reached around 0.9, isopropyl β-D-1-thiogalactopyranoside (IPTG) was added to 0.5 mM to induce recombinant protein expression. After cultivating overnight at 20 °C and 110 rpm, *E. coli* cells were harvested by centrifugation at 7000×g at 4 °C for 10 min, then suspended in binding buffer (20 mM Tris, 500 mM NaCl, 5 mM imidazole, pH 8.0). After the sonication for 16 min, the lysate was centrifuged at 40,000×g, 4 °C for 30 min. The supernatant was purified on a Ni-NTA agarose column (QIAGEN, Germany), pre-equilibrated with the binding buffer. The His-tagged protein was eluted with elution buffer (20 mM Tris, 500 mM NaCl, 200 mM imidazole, pH 8.0). The protein was then purified over a Hiload 16/600 Superdex 75pg column (Cytiva, Marlborough, Massachusetts) equilibrated with PBS buffer.

### Cloning, expression, and purification of HigB2

The preparation of HigB2 was carried out as described previously with several changes ^49, 50^. Briefly, the expression-optimized genes encoding HigB2 and HigA2 were inserted into pET-Duet vector (Novagen, Madison, Wisconsin), and a His_6_-tag followed by thrombin cleavage site was introduced into N-terminal of HigB2. *E.coli* strain of C43(DE3) was transformed with the expression vectors, then cultivated in 1 L of LB medium containing 100 µg/mL ampicillin at 37 °C and 125 rpm. Expression of the HigB2–HigA2 complex was induced by adding 0.5 mM IPTG when the optical density at 600 nm reached 0.6. After cultivating overnight at 20 °C and 100 rpm, *E. coli* cells were harvested by centrifugation at 7000×g at 4 °C for 10 min, then suspended in lysis buffer (50 mM Tris, 200 mM NaCl, 1 mM PMSF, pH 8.0). After the sonication and centrifugation, the supernatant was added to a 1-mL Ni-NTA agarose column (QIAGEN). The column-bound HigB2–HigA2 complex was disrupted by washing the column with buffer containing 5 M guanidine-HCl, and then a simple refolding process, HigB2 was eluted with buffer containing 300 mM imidazole. The protein was then purified over a Hiload 16/600 Superdex 75pg column (Cytiva) equilibrated with 20 mM Tris, 200 mM NaCl, pH 8.0.

### Circular dichroism (CD) measurements

Far ultraviolet Circular Dichroism (CD) spectra were measured on a J-820 spectropolarimeter (Jasco, Japan). Measurements were performed in 1-mm quartz cuvettes in PBS buffer at a concentration of 8 µM for antibodies or 7 µM for HigB2. Each spectrum shown is an accumulation of five measurements.

### Surface plasmon resonance (SPR) analysis

Interactions between the antigen and the antibodies were analyzed in a Biacore T200 instrument (Cytiva). HigB2 (4 µg/mL) in 10 mM sodium acetate at pH 5.5 was immobilized onto a CM5 Sensor Chip at 130 RU by the amine-coupling method. Measurements were carried out in PBS buffer supplemented with 0.005% (v/v) Tween-20. Binding of antibodies were determined by increasing the concentration of the analyte at a flow rate of 30 µL/min at 25 °C. Both of the association and dissociation times were 120 s or 180 s. Antibodies were subsequently dissociated from the ligand by regeneration buffer (10 mM Glycine-HCl pH 2.0). Kinetic parameters (*k*_on_, *k*_off_, *K*_D_) were calculated with the BIAevaluation software (Cytiva) using a 1:1 global fitting model.

### Differential scanning calorimetry (DSC) measurements

Thermal stability of Nb8 and mutants (70 µM) in PBS buffer was monitored with a MicroCal PEAQ-DSC instrument (Malvern Panalytical, United Kingdom). Samples were scanned at a speed of 1 °C /min from 20 to 110 °C. Data analysis was performed with MicroCal PEAQ-DSC software (Malvern Panalytical) using a non-two-state denaturation model.

### Molecular dynamics (MD) simulations

MD simulations of the Nb8 monomer were performed using GROMACS 2018.6 ^51^ with the CHARMM36m ^52^ force field. Chain B of the 5mje co-crystal was used as the initial structure. Using the CHARMM-GUI ^53^, the initial structures were solvated with TIP3P water ^54^ in a rectangular box such that the minimum distance to the edge of the box was 15 Å under periodic boundary conditions. Sodium and chloride ions were added to neutralize protein charge, then additional ions were added to a salt concentration of 0.15 M. The time step was set to 2 fs throughout the simulations. Energy minimization was performed and equilibration was conducted with the NVT ensemble (298 K) for 1 ns. Further simulations were performed with the NPT ensemble at 298 K. Structural snapshots were saved every 10 ps. Distances between two aromatic ring centroids or two atoms were calculated using GROMACS commands. Distance cutoffs were set at 6.5 Å for π-π interaction (centroid-centroid distance) ^55, 56^, 3.5 Å for hydrogen bonds (donor to acceptor distance) ^56–58^, and 4.0 Å for salt bridges (N-O distances) ^56, 59, 60^.

### Hydrogen/deuterium exchange mass spectrometry (HDX-MS) measurements

WT Nb8 and Y93A, E94A, and E95A mutants were prepared in PBS at final concentrations of 2.0 mg/mL. Each protein was diluted 10-fold with PBS in D_2_O. The diluted solutions were then incubated at 10 °C. Deuterium-labeled samples were quenched by diluting by about two folds with quenching buffer composed of 8 M urea and 1 M Tris (2-carboxyethyl) phosphine hydrochloride at pH 3.0. All dilutions were performed using an HDx-3 PAL (LEAP Technologies, Morrisville, North Carolina). After quenching, the solutions were subjected to online pepsin digestion followed by LC/MS analysis using an UltiMate3000RSLCnano (Thermo Fisher Scientific, Waltham, Massachusetts) connected to a Q Exactive plus mass spectrometer (Thermo Fisher Scientific). Online pepsin digestion was performed on a protease type XIII /pepsin column (w/w, 1:1; 2.1 × 30 mm; NovaBioAssays Inc., Woburn, Massachusetts) in formic acid solution (pH 2.5) for 3 min at 8 °C at a flow rate of 50 μL/min. After pepsin digestion desalting and analysis were performed using Acclaim PepMap 300 C18 (1.0 × 15 mm, Thermo Fisher Scientific) and Hypersil GOLD (1.0 × 50 mm, Thermo Fisher Scientific) columns. The mobile phase was 0.1% formic acid solution (buffer A) and 0.1% formic acid containing 90% acetonitrile (B buffer). The deuterated peptides were eluted at a flow rate of 45 μL/min with a gradient of 10% to 90% of buffer B for 9 min. The conditions of the mass spectrometer were as follows: electrospray voltage, 3.8 kV; positive ion mode; sheath and auxiliary nitrogen flow rates at 20 and 2 arbitrary units, respectively; ion transfer tube temperature, 275 °C; auxiliary gas heater temperature, 100 °C; and mass range, 200 to 2000 m/z. Data-dependent acquisition was performed using a normalized collision energy of 27 arbitrary units. The MS and MS/MS spectra were subjected to a database search analysis using the Proteome Discoverer 2.2 (Thermo Fisher Scientific). Analysis of the deuteration levels of the peptide fragments was performed by comparing the spectra of deuterated samples with those of nondeuterated samples using the HDExaminer software (Sierra Analytics, Modesto, California). Data summary for each HDX-MS is shown (Table S3).

## Acknowledgements

This work was funded in part by Japan Society for the Promotion of Science under grant number JP19H05766, JP20H02531 (to K.T.), Japan Agency for Medical Research and Development under grant number JP22ama121033j (to K. T.), and the JST CREST under grant number JPMJCR20H8 (to K. T.). This research was partially supported by Research Support Project for Life Science and Drug Discovery (Basis for Supporting Innovative Drug Discovery and Life Science Research (BINDS)) from AMED under Grant Number 22ama121033 (to K. T.). The supercomputing resource in this study was provided by the Human Genome Center at the Institute of Medical Science, The University of Tokyo.

## Notes

### Competing Interest Statement

The authors have declared no competing interest.

