## Supporting Information for "Conformational features and interaction mechanisms of VHH antibodies with β-hairpin-like CDR-H3: A case of Nb8-HigB2 interaction"

**Table S1.** Residues at the Nb8-HigB2 interface.

| **Position** | | **Contact interface** | | |
|---|---|---|---|---|
| **Region** | **Chothia** | **BSA[a] (Å$^2$)** | **BSA/ASA[a] (%)** | **Polar contacts[b]** |
| **FR1** | Q1 | 28.3 | 18.7 | |
| | L4 | 24.8 | 52.8 | h |
| **FR2** | F37[c] | 51.2 | 86.5 | |
| | Q39 | 51.7 | 57.3 | |
| | P41 | 60.0 | 51.3 | |
| | Q44 | 34.4 | 20.9 | |
| | L47 | 64.9 | 75.3 | |
| | T50 | 13.0 | 52.0 | H |
| **FR3** | F89 | 71.3 | 76.7 | |
| | Y93[c] | 22.4 | 90.0 | |
| | (E94[c]) | 0.00 | 0.0 | |
| **CDR3** | E95[c] | 39.0 | 89.4 | H, S |
| | R97[c] | 26.7 | 31.1 | H, S |
| | S100b[c] | 33.1 | 45.3 | H |
| | R100c[c] | 116 | 74.8 | h, H, S |
| | N100d[c] | 35.5 | 78.7 | |
| | T101[c] | 64.3 | 99.2 | h, H |
| | Y102[c] | 23.9 | 56.6 | |
| **FR4** | W103[c] | 118 | 91.5 | h |
| | Q105 | 79.9 | 46.9 | |
| | Q108 | 47.3 | 62.4 | |
| | T110 | 20.4 | 58.4 | |

[a] BSA, buried surface area; ASA, accessible surface area.

[b] h, hydrogen bond (main chain); H, hydrogen bond (side chain); S, salt bridge.

[c] Residues mutated in alanine-scanning experiments.

**Table S2.** Kinetic parameters of wild-type Nb8 and mutants determined using SPR. Error bars correspond to the standard deviation. *N.D.*: not determined.

| | $k_{on}$ ($10^{5}$ $M^{-1}$ $s^{-1}$) | $k_{off}$ ($10^{-3}$ $s^{-1}$) | $K_D$ ($10^{-8}$ M) |
|---|---|---|---|
| **WT** | 2.32±0.03 | 3.95±0.05 | 1.70±0.04 |
| **F37A** | 1.97±0.09 | 12.6±1.6 | 6.47±1.11 |
| **Y93A** | *N.D.* | *N.D.* | *N.D.* (>1600) |
| **E94A** | 0.300±0.080 | 161±47 | 536±12 |
| **E95A** | 7.52±2.43 | 10.4±3.6 | 1.37±0.05 |
| **R97A** | 0.197±0.019 | 59.4±4.1 | 302±9 |
| **S110bA** | 1.81±0.03 | 7.86±0.08 | 4.35±0.05 |
| **R100cA** | 0.303±0.020 | 167±11 | 550±4 |
| **N110dA** | 3.41±0.03 | 3.88±0.11 | 1.14±0.03 |
| **T101A** | 3.04±0.24 | 21.2±0.3 | 7.01±0.48 |
| **Y102A** | 2.67±0.05 | 5.54±0.06 | 2.08±0.04 |
| **W103A** | 0.274±0.004 | 34.9±0.2 | 127±2 |

**Table S3.** Summary of HDX-MS experiments

(A) Y93A

| Data Set | WT (control) | Y93A |
|---|---|---|
| HDX time course (sec) | 60, 120, 240, 480, 960, 1920, 3840 | 60, 120, 240, 480, 960, 1920, 3840 |
| # of peptides | 264 | 252 |
| Sequence coverage | 100.00% | 96.92% |
| Average peptide length / Redundancy | 16.37 / 33.24 | 16.02 / 31.06 |
| Replicates | 2 | 2 |
| Repeatability (avg. stddev of #D) | 0.1631 | 0.1893 |

(B) E94A

| Data Set | WT (control) | E94A |
|---|---|---|
| HDX time course (sec) | 60, 120, 240, 480, 960, 1920, 3840 | 60, 120, 240, 480, 960, 1920, 3840 |
| # of peptides | 261 | 251 |
| Sequence coverage | 100.00% | 100.00% |
| Average peptide length / Redundancy | 16.25 / 32.62 | 15.78 / 30.48 |
| Replicates | 2 | 2 |
| Repeatability (avg. stddev of #D) | 0.1574 | 0.1416 |

(C) E95A

| Data Set | WT (control) | E95A |
|---|---|---|
| HDX time course (sec) | 60, 120, 240, 480, 960, 1920, 3840 | 60, 120, 240, 480, 960, 1920, 3840 |
| # of peptides | 264 | 225 |
| Sequence coverage | 100.00% | 97.69% |

| | | |
|---|---|---|
| Average peptide length / Redundancy | 16.37 / 33.24 | 16.77 / 29.02 |
| Replicates | 2 | 2 |
| Repeatability (avg. stddev of #D) | 0.1631 | 0.2355 |

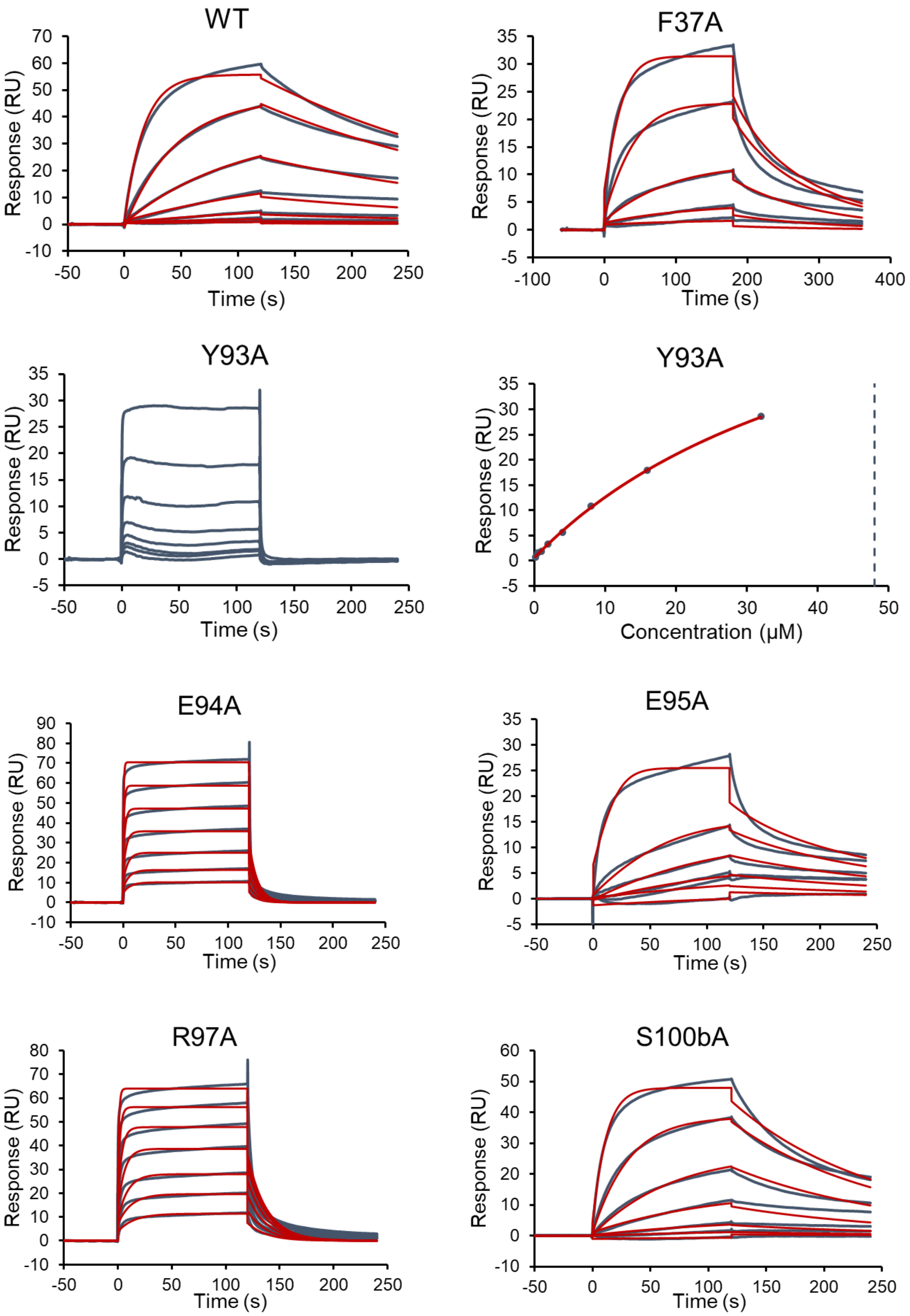
WT
Response (RU)
Time (s)
F37A
Response (RU)
Time (s)
Y93A
Response (RU)
Time (s)
Y93A
Response (RU)
Concentration (μM)
E94A
Response (RU)
Time (s)
E95A
Response (RU)
Time (s)
R97A
Response (RU)
Time (s)
S100bA
Response (RU)
Time (s)

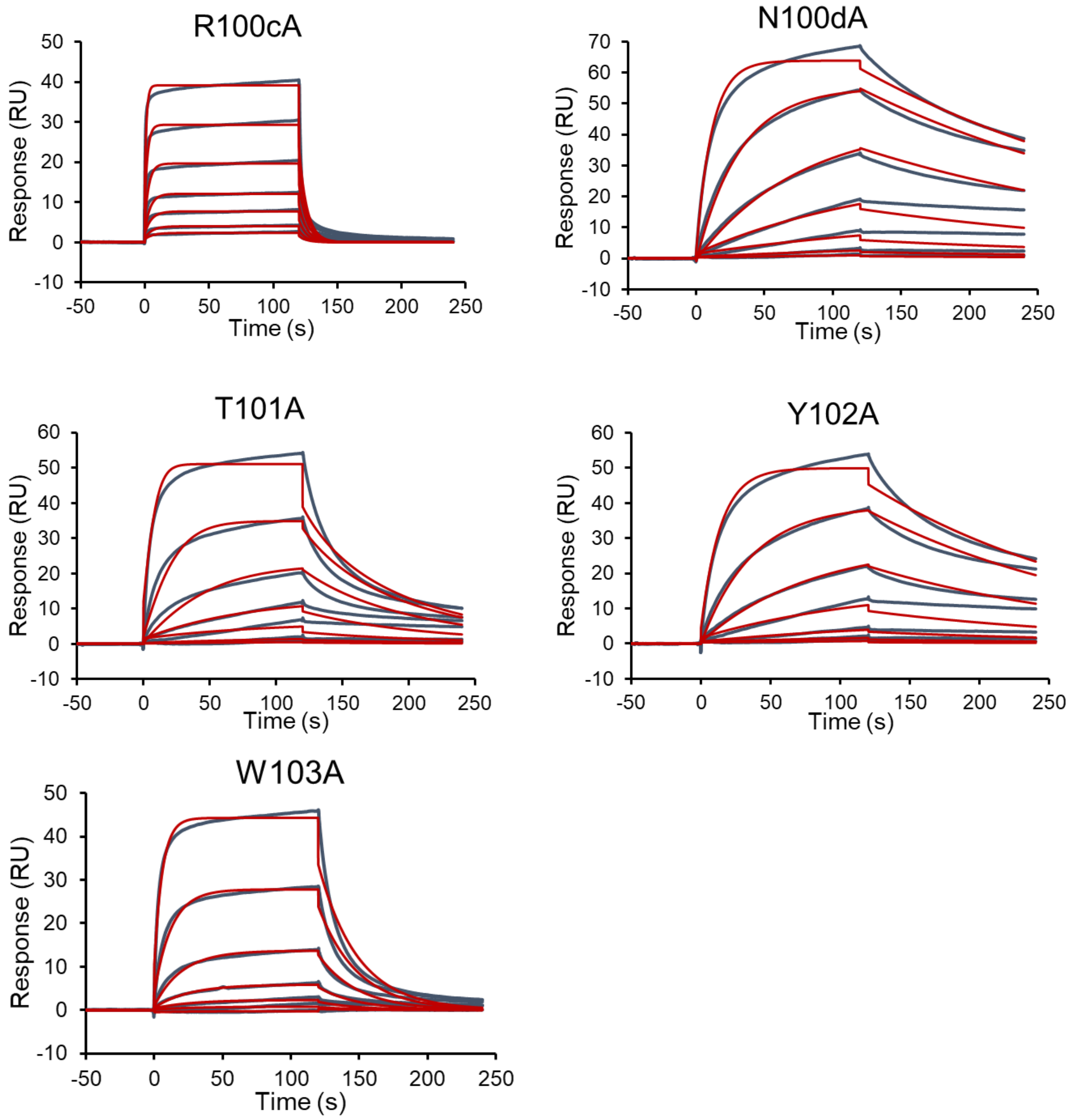


**Figure S1.** SPR sensorgrams of the binding of Nb8 to HigB2 at 25 °C. Due to high association and dissociation rates, the Scatchard plot for Y93A was depicted based on maximum steady-state responses at each concentration. Concentration ranges were 243~0.33 nM for WT, N100dA, and Y102A; 320, 160~2.5 nM for F37A; 32~0.25 µM for Y93A; 64~1 µM for E94A; 128, 32~2 nM for E95A; 48~0.75 µM for R97A; 486~0.67 nM for S100bA; 19.2~0.3 µM for R100cA; 729~1 nM for T101A; and 4860~6.67 nM for W103A.

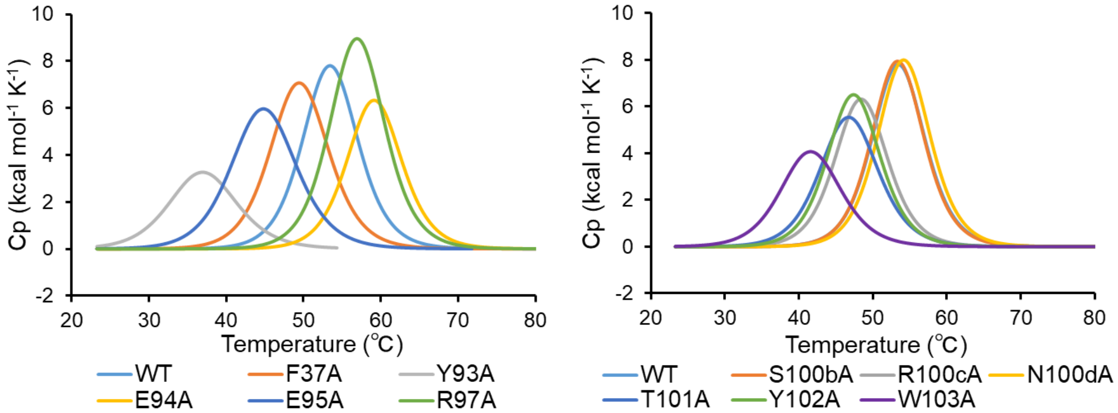


**Figure S2.** DSC thermograms of Nb8 antibodies. Acquired data were fit to a non-two-state model.

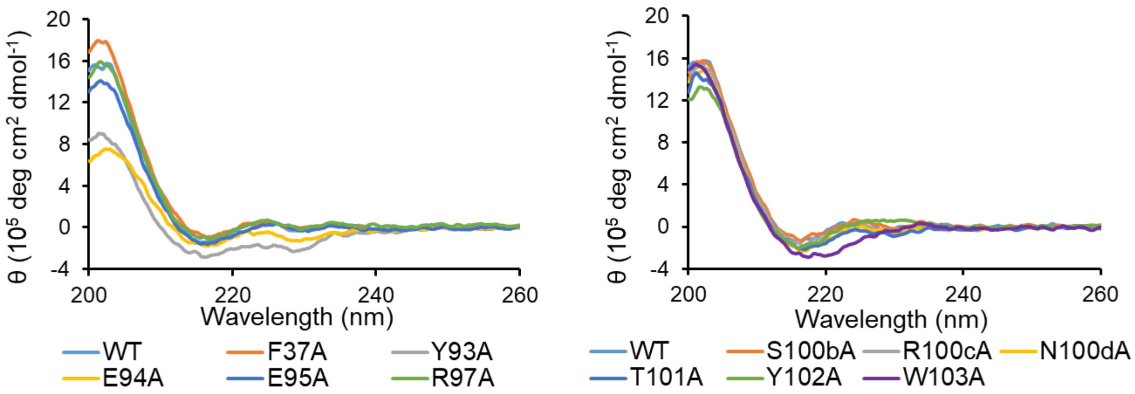


**Figure S3.** CD spectra of Nb8 antibodies. Each spectrum is the average of five measurements.

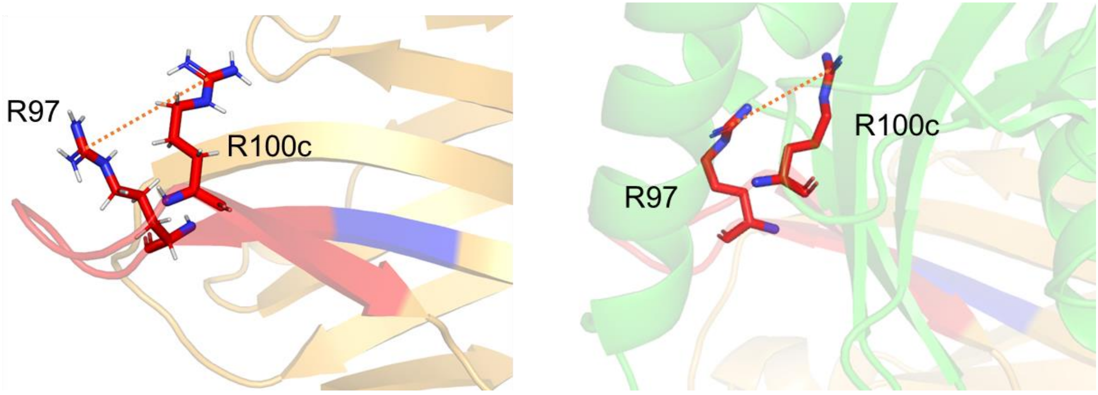


**Figure S4.** Adjacent locations and electric repulsions between R97 and R100c in MD simulation (left; run 1, 900.00 ns) and co-crystal structure (right).

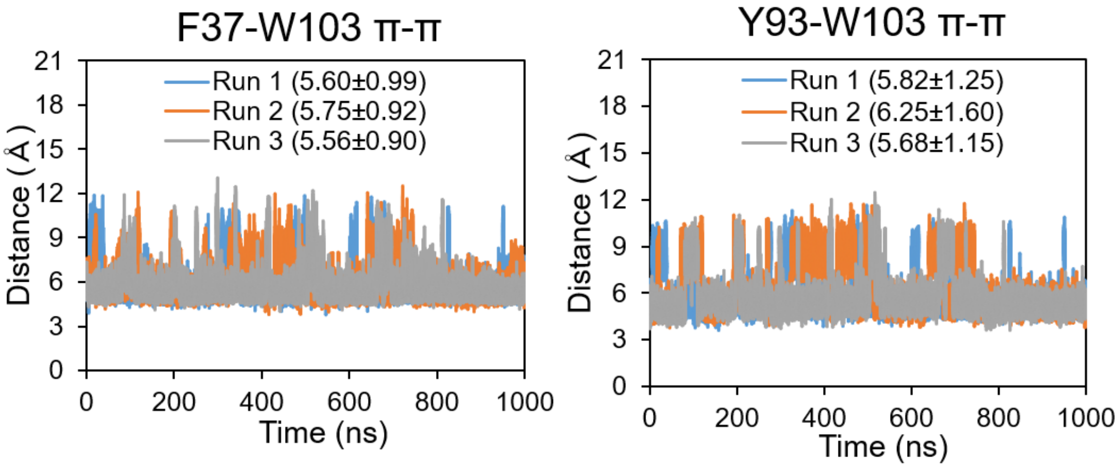


**Figure S5.** Distance fluctuations of π-π interactions. Centroids of each aromatic ring were used for the analysis. Averages and standard deviations of distances are given.

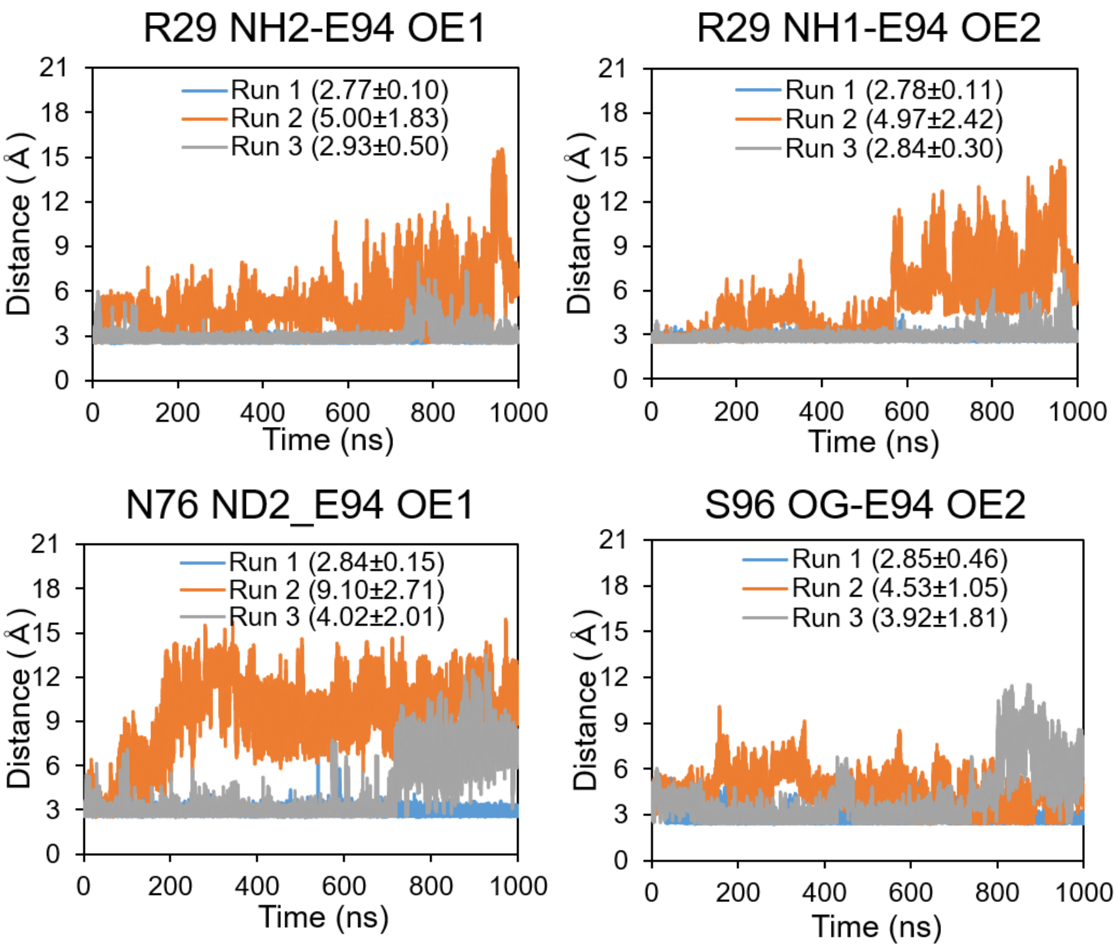


**Figure S6.** Changes in distances in intramolecular hydrogen bond networks over simulation time. Averages and standard deviations are given.

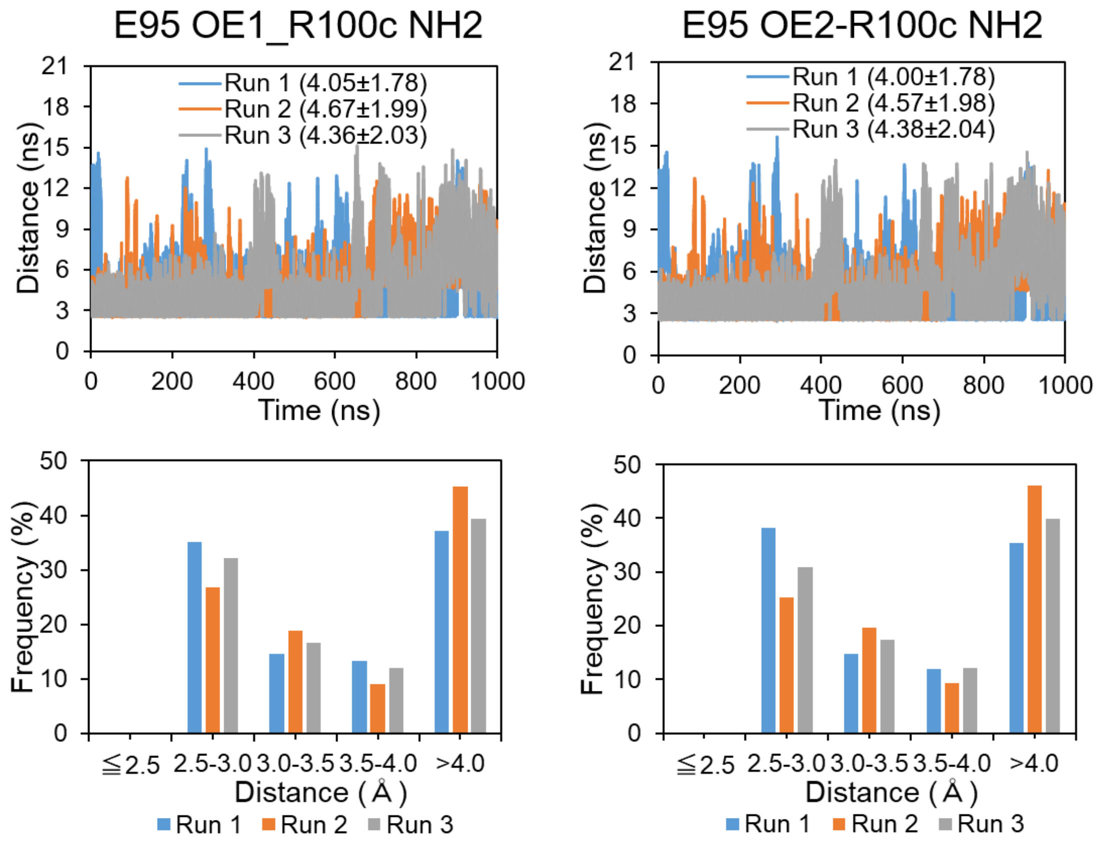


**Figure S7.** Distance changes and distribution in E95-R100cA interactions. Averages and standard deviations are given.

(A)

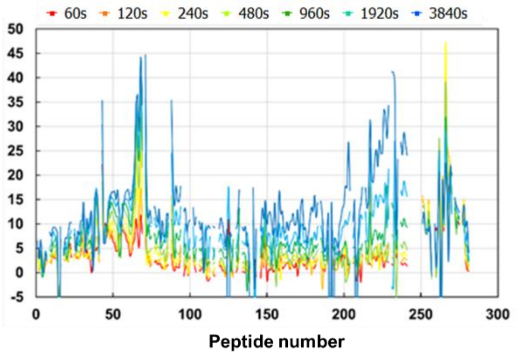
60s 120s 240s 480s 960s 1920s 3840s
D (%)
Peptide number

(B)

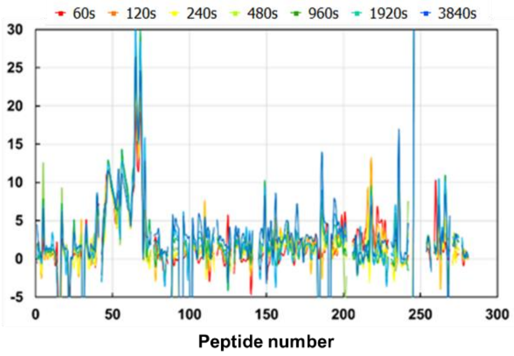
60s 120s 240s 480s 960s 1920s 3840s
D (%)
Peptide number

(C)

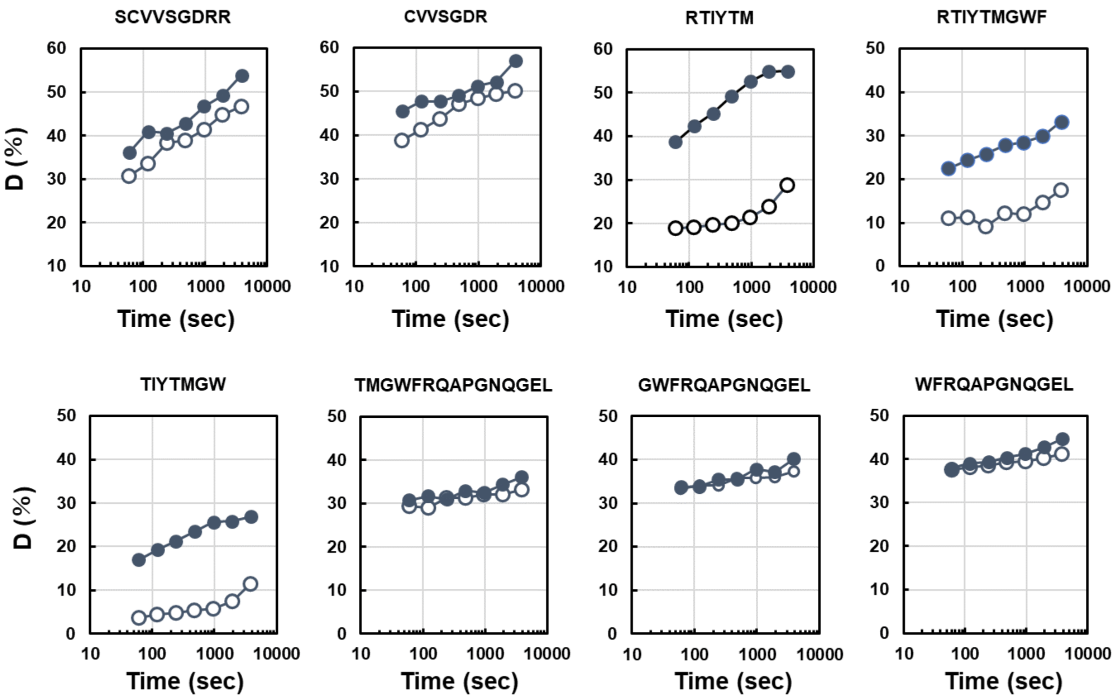
SCVVSGDRR
CVVSGDR
RTIYTM
RTIYTMGWF
TIYTMGW
TMGWFRQAPGNQGEL
GWFRQAPGNQGEL
WFRQAPGNQGEL
D (%)
Time (sec)

(D)

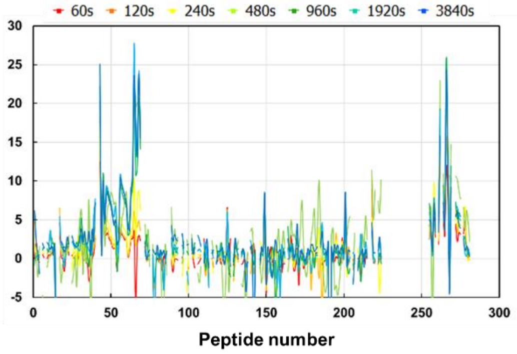
60s 120s 240s 480s 960s 1920s 3840s
D (%)
Peptide number

**(E)**

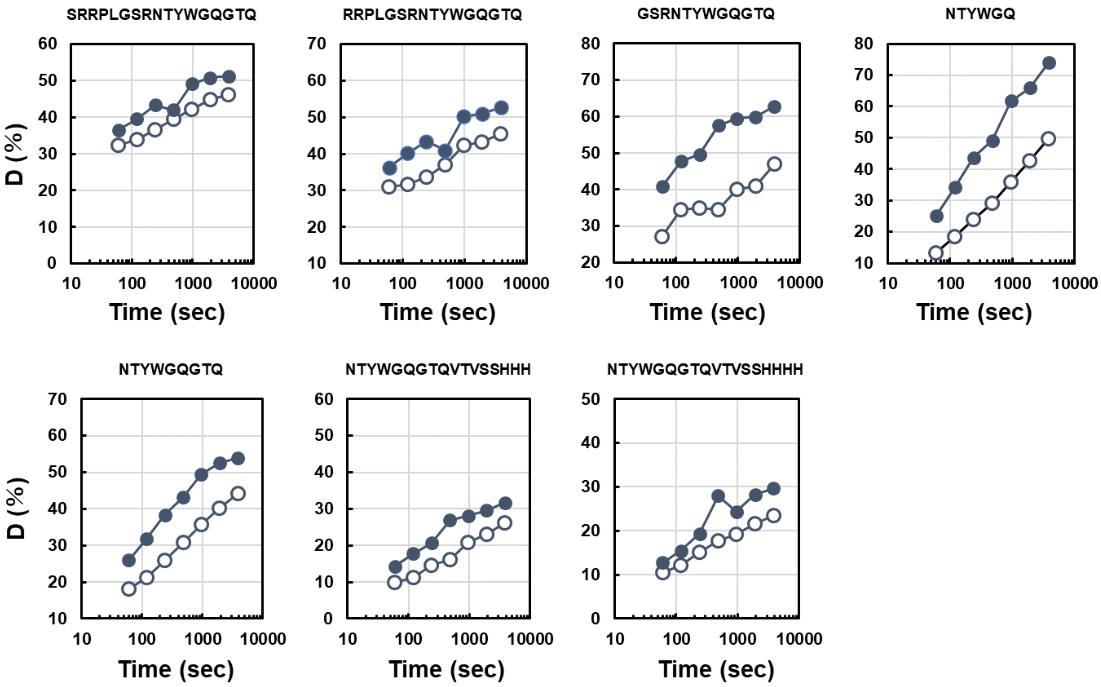


**Figure S8.** HDX-MS analysis of Nb8 and mutants. (A) Differences of HDX rates between Nb8 WT and Y93A. (B) Differences of HDX rates between Nb8 WT and E94A. (C) HDX-MS profiles of indicate regions of Nb8 WT and E94A. (D) Differences of HDX rates between Nb8 WT and E95A. (E) HDX-MS profiles of indicate regions of Nb8 WT and E95A.

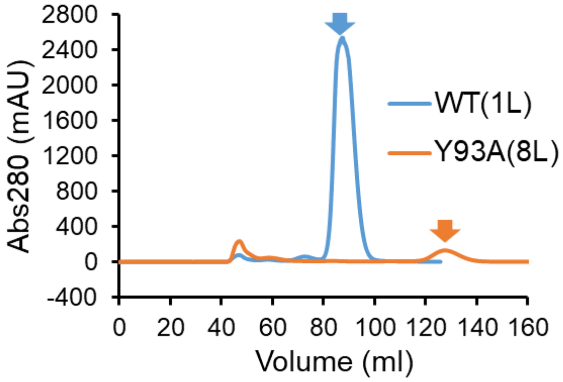


**Figure S9.** SEC chromatogram of Nb8 WT and Y93A mutant. Corresponding peaks of target proteins are indicated by arrows. Total cultivation volumes in *E.coli* expression step are also depicted.

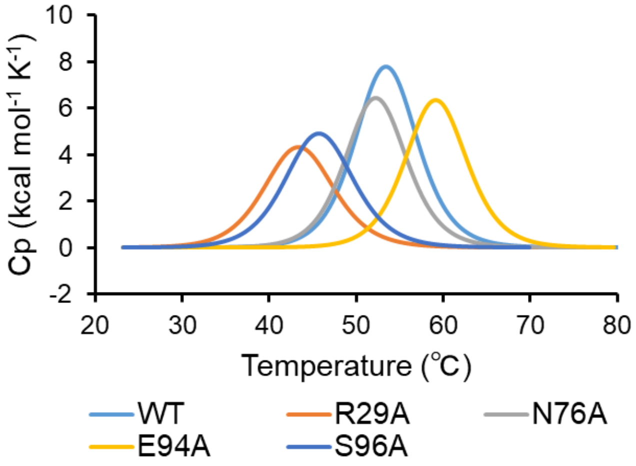


| | $T_m$ (°C) | ΔH (kcal/mol) |
|---|---|---|
| **WT** | 53.5±0.0 | 73.4±1.7 |
| **R29A** | 43.4±0.1 | 45.0±1.3 |
| **N76A** | 52.2±0.2 | 60.2±0.5 |
| **E94A** | 59.1±0.1 | 59.1±2.4 |
| **S96A** | 45.4±0.3 | 49.7±2.0 |

**Figure S10.** DSC thermograms of Nb8 mutants involved in buried hydrogen bond network. Acquired data was fitted by non-two-state model. Thermostability parameters are given in lower panel with standard deviations.

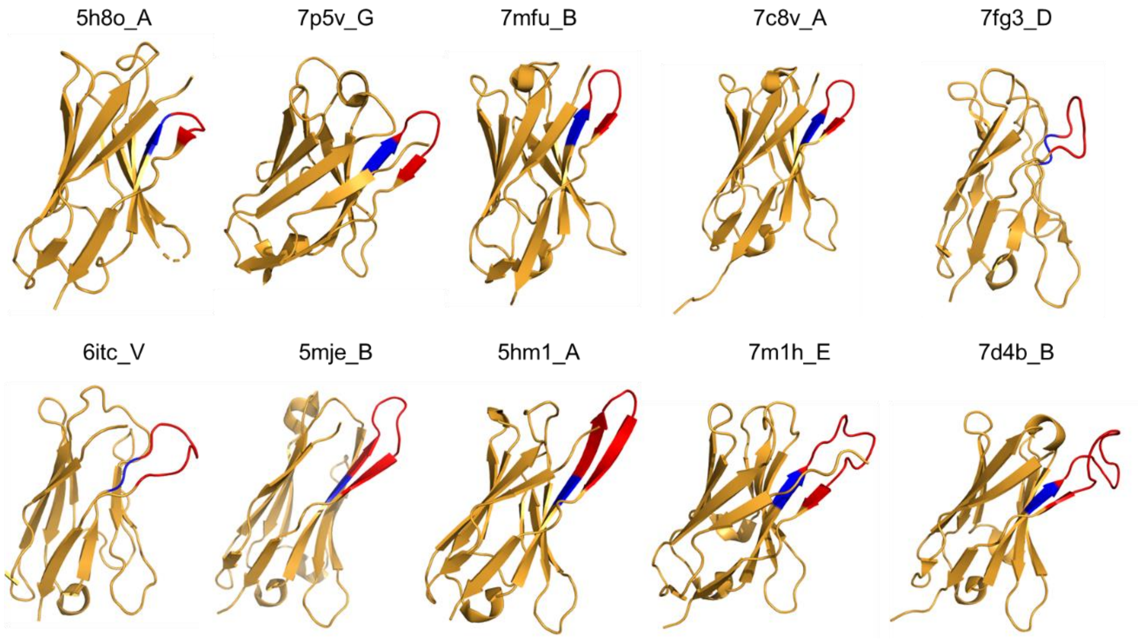


**Figure S11.** Crystal structures of VHH antibodies harboring Y93. Antigens aren't shown in this figure. Only the CDR3 of 7d4b_B isn't regarded as β-hairpin-like conformation.
